# Clinical and immunological signatures of severe COVID-19 in previously healthy patients with clonal hematopoiesis

**DOI:** 10.1101/2021.10.05.463271

**Authors:** Chang Kyung Kang, Baekgyu Choi, Sugyeong Kim, Seongwan Park, Soon Ho Yoon, Dohoon Lee, Andrew J. Lee, Yuji Ko, Euijin Chang, Jongtak Jung, Pyoeng Gyun Choe, Wan Beom Park, Eu Suk Kim, Hong Bin Kim, Nam Joong Kim, Myoung-don Oh, Suk-jo Kang, Kyuho Kang, Sun Kim, Hogune Im, Joohae Kim, Yong Hoon Lee, Jaehee Lee, Ji Yeon Lee, Joon Ho Moon, Kyoung-Ho Song, Youngil Koh, Inkyung Jung

## Abstract

Identifying additional risk factors for COVID-19 severity in numerous previously healthy patients without canonical clinical risk factors remains challenging. In this study, we investigate whether clonal hematopoiesis of indeterminate potential (CHIP), a common aging-related process that predisposes various inflammatory responses, may exert COVID-19 severity. We examine the clinical impact of CHIP in 143 laboratory-confirmed COVID-19 patients. Both stratified analyses and logistic regression including the interaction between canonical risk factors and CHIP show that CHIP is an independent risk factor for severe COVID-19, especially in previously healthy patients. Analyses of 60,310 single-cell immune transcriptome profiles identify distinct immunological signatures for CHIP (+) severe COVID-19 patients, particularly in classical monocytes, with a marked increase in pro-inflammatory cytokine responses and potent IFN-γ mediated hyperinflammation signature. We further demonstrate that the enhanced expression of CHIP (+) specific IFN-γ response genes is attributed to the CHIP mutation-dependent epigenetic reprogramming of poised or bivalent *cis*-regulatory elements. Our results highlight a unique immunopathogenic mechanism of CHIP in the progression of severe COVID-19, which could be extended to elucidate how CHIP contributes to a variety of human infectious diseases.

## Introduction

A pandemic of coronavirus disease-19 (COVID-19), an emerging infectious disease caused by severe acute respiratory syndrome-coronavirus-2 (SARS-CoV-2), has been being a global health threat of the century^1,2^. Epidemiologic data revealed that about 20% of patients underwent severe or critical course^3^. Although many clinical risk factors for severe illness including older age, comorbidities such as diabetes mellitus or hypertension, and morbid obesity have been found^4,5^, we could still only partly explain the development of severe COVID-19. Indeed, there have been numerous cases of severe COVID-19 from previously healthy adults^6,7^.

Clinical deteriorations such as acute respiratory distress syndrome or intensive care unit admission most commonly occur at around the 10^th^ day of illness^8,9^, when the viral loads are declining after the early peak^10,11^. This temporal discrepancy suggests immunological phenomenon may play an important role in the progression of COVID-19. High levels of circulating pro-inflammatory cytokines^12^, aberrant hyperactivation of cytotoxic lymphocytes^13^ or their infiltration in vital organs^14^, or dysregulated monocytes and macrophages^15^ have been proposed as mechanisms for pathologic immune responses in severe COVID-19.

Clonal hematopoiesis of indeterminate potential (CHIP) refers to a population of immune cells with acquired gene mutations, but without fulfilling diagnostic criteria for a hematologic malignancy^16^. As a majority of the genes associated with CHIP including *DNMT3A, TET2*, and *ASXL1* are involved in epigenetic regulation, CHIP may have a wide range of effects on immune function through altered chromatin activities^17^. There is growing evidence supporting a role for the CHIP mutations in altered immune function through effector cells such as monocytes/macrophages and their dysregulated cytokine/chemokine expression, which account for the increased risk of cardiovascular disease in individuals with CHIP^18-21^. Since immunopathogenesis of such an adverse outcome of CHIP largely shares that of severe COVID-19, we hypothesized that CHIP might contribute to the progression of COVID-19 with its unique immune signature. Although recent reports describe the association between acquired mutations in hematopoietic cells such as CH and severe COVID-19^22,23^, there has been a controversy on the clinical impact of CHIP in COVID-19^23-25^. However, if there exists a distinct CHIP-related clinical deterioration mechanism of COVID-19, tailored therapeutics could be of help to salvage these patients.

In this study, to determine the clinical significance of CHIP in COVID-19 severity, especially in previously healthy adults, we analyzed thorough clinical, radiological, and laboratory characteristics of patients with COVID-19. In addition, we explored immune signatures using single-cell RNA expression data according to the presence of CHIP to suggest how CHIP attributes to the immunologic responses in severe COVID-19. Lastly, since the majority of mutations in CHIP are related to epigenetic regulators of DNA methylation and heterochromatin formation, we investigated CHIP mutation dependent dysregulated epigenetic gene regulation mechanisms involved in CHIP-specific immunopathogenesis.

## Results

### Impact of CHIP on severe COVID-19 in previously healthy patients

A total of 143 laboratory-confirmed COVID-19 patients were analyzed in this study (Fig. 1a). Among those, 34 patients had CHIP (23.8%, Supplementary Table 1). *DNMT3A* (13 variants) was the most common mutated gene, followed by *TET2* (7 variants) and *ASXL1* (5 variants). Clinical characteristics were retrospectively reviewed using electronic medical record (EMR) systems of each institution. It included age, sex, body mass index, presence of comorbidities, details of oxygen and medical therapy, duration of hospital stay, and in-hospital mortality. Serial laboratory findings including complete blood count with differential count and chemistries were also collected. Cases with the highest ordinal scale 3 (need for supplemental oxygen therapy via nasal cannula) or more were classified as severe COVID-19, while others were classified with mild ones^3^.

**Figure 1.**
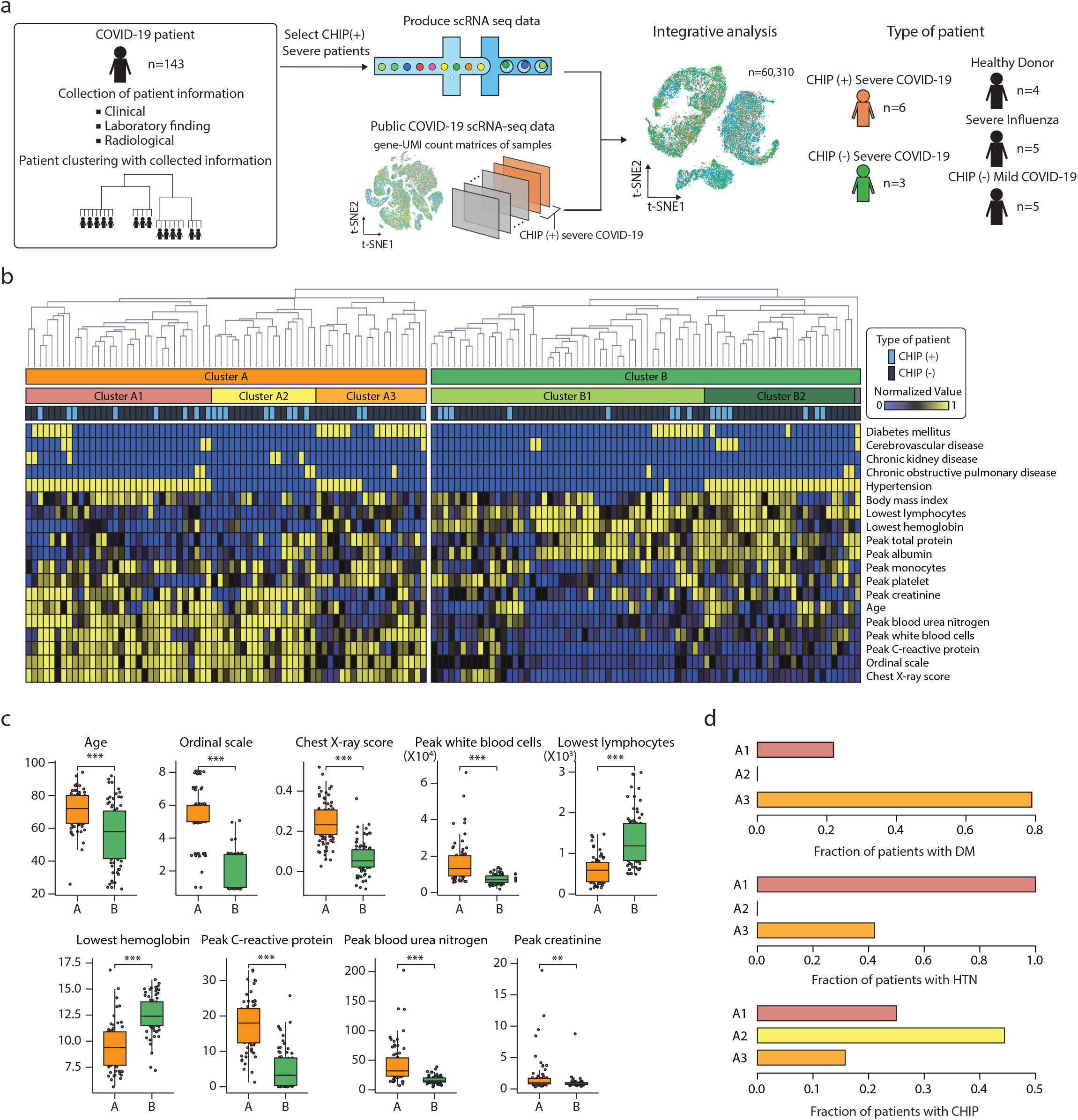
Hierarchical clustering analysis for clinical characteristics of severe COVID-19. **a**, Overview of study design. **b**, A heatmap shows the hierarchical clustering result of clinical characteristics. On the top, three different bars represent clusters and the presence of CHIP mutations for each patient. The first bar represents two main clusters of patients (n=69 for cluster A and n=74 for cluster B). The second bar represents five sub-clusters of patients (n=32 for A1, n=18 for A2, n=19 for A3, n=47 for B1, and n=26 for B2). The third bar represents the presence of CHIP. **c**, Boxplots showing clinical characteristics between cluster A (orange) and B (green). For the boxplots, the box represents the interquartile range (IQR) and the whiskers correspond to the highest and lowest points within 1.5 × IQR. Statistical significance was examined with unpaired t-test (* < p-value 0.05, ** < p-value 0.01, and *** < p-value 0.001). **d**, Barplots showing fractions of patients with comorbidities or CHIP for cluster A1, A2, and A3. HTN stands for hypertension. DM stands for diabetes mellitus.

Baseline characteristics of these patients are shown in Supplementary Table 1. Median (interquartile range [IQR]) ages were 73 (61—81) and 65 (50—75) years in those with or without CHIP, respectively (Student’s *t-*test, *P* < 0.001), as consistent with a common aging-related property of CHIP. The presence of CHIP appears to contribute to COVID-19 severity as severe COVID-19 tended to be more frequent in patients with CHIP than those without, while the difference was statistically not significant (25/34, 73.5% vs. 65/109, 59.6%, Chi-squared test, *P* = 0.143).

To precisely examine the clinical impact of CHIP in patients with COVID-19, we conducted a hierarchical clustering analysis for baseline characteristics and examined the distribution of CHIP among clusters (Fig. 1b). All continuous clinical information was transformed into a range of 0 to 1 using a logistic function, and comorbidity status was dichotomized into 0 and 1 for absence and presence (see Methods). Patients were grouped into two main clusters, A and B. They mainly comprised severe and mild cases, respectively, since cluster A had significantly higher ordinal scores, peak serum C-reactive protein level, and peak chest X-ray score than cluster B (Unpaired t-test, *P* < 0.001) (Fig. 1c). Cluster A was subdivided into three clusters, A1, A2, and A3 with their own comorbidity status (Fig. 1d). Cluster A1 was characterized by the presence of hypertension, while cluster A3 was a DM-enriched group.

Interestingly, cluster A2 tended to have a higher rate of CHIP than the others in cluster A (Fisher’s exact test, *P* = 0.074), while it had significantly lower BMI (median [IQR], 20.8 [17.8—23.1] vs. 23.0 [21.7—26.1]; Student’s *t-*test, *P* = 0.005) and lower rate of DM (0/18, 0% vs. 22/51, 43.1%; Fisher’s exact test, *P* = 0.001) or hypertension (0/18, 0% vs. 40/51, 78.4%; Fisher’s exact test, *P* < 0.001) (Fig. 1d and Supplementary Table 2). However, age distribution was not statistically different (Fisher’s exact test, *P* = 0.662).

The presence of a particular type of severe cases uniquely enriched by CHIP without canonical risk factors such as DM, hypertension, and high BMI led us to hypothesize that CHIP may contribute to severe COVID-19 in its own way. An additional hierarchical clustering with DM, hypertension, and CHIP information in all severe patients (Ordinal scale >=3) also supported our hypothesis since it showed that CHIP was clustered together (Extended Data Fig. 1).

To test the statistical significance of the impact of CHIP on COVID-19 severity in previously healthy patients, we stratified the entire patients by the presence of any of the canonical risk factors such as DM, hypertension, and BMI ≥ 30.0. When adjusted with age and gender, the risk of severe COVID-19 was significantly higher in CHIP (+) patients than in CHIP (−) ones in canonical risk factor-absent subgroup (adjusted odds ratio [95% confidence interval], 14.8 [1.3—164.1]; logistic regression *P* = 0.028; Table 1). Multivariate analysis including the interaction between canonical risk factors and CHIP revealed that CHIP was an independent risk factor for severe COVID-19 (adjusted odds ratio [95% confidence interval], 10.7 [1.1—100.7], logistic regression *P* = 0.038, Table 2). The results of the patients clustering and the statistical evidence indicate that CHIP is an independent risk factor for severe COVID-19, especially in previously healthy patients.

**Table 1.** Age and gender-adjusted odds ratio of the presence of CHIP for severe COVID-19 in canonical risk factors-stratified subgroups.

|  | Severe COVID-19 |  | aOR (95% CI) | *P* |
| --- | --- | --- | --- | --- |
|  | CHIP (+) | CHIP (-) |  |  |
| Canonical risk factor present | 13/21 (61.9%) | 47/68 (69.1%) | 0.4 (0.1—1.4) | 0.159 |
| Canonical risk factor absent | 12/13 (92.3%) | 18/41 (43.9%) | 14.8 (1.3—164.1) | 0.028 |
| Total | 25/34 (73.5%) | 65/109 (59.6%) | 1.1 (0.4—2.8) | 0.829 |

CHIP, clonal hematopoiesis with indeterminate potential; aOR, adjusted odds ratio; CI, confidence interval
Canonical risk factors were BMI ≥ 30.0, diabetes mellitus, and hypertension.

**Table 2.** Final multivariate model for independent risk factors for severe COVID-19 including interaction term between CHIP and canonical risk factors.

|  | aOR | *P* |
| --- | --- | --- |
| Age, per one year | 1.1 (1.0—1.1) | < 0.001 |
| Male gender | 2.0 (0.9—4.5) | 0.075 |
| CHIP | 10.7 (1.1—100.7) | 0.038 |
| Canonical risk factors | 2.0 (0.8—5.0) | 0.120 |
| CHIP * conventional risk factors | 0.0 (0.0—0.5) | 0.012 |

CHIP, clonal hematopoiesis with indeterminate potential; aOR, adjusted odds ratio
Canonical risk factors were BMI ≥ 30.0, diabetes mellitus, and hypertension.

### Distinct immune signatures in CHIP (+) severe COVID-19

We next sought to identify distinct immune signatures for severe COVID-19 according to the presence of CHIP. To this end, a total of 60,310 high-quality single-cell transcriptome profiles of peripheral blood mononuclear cells (PBMCs) generated by 10x Genomics single-cell RNA-seq (scRNA-seq) platform were integrated from healthy donors (n=4), severe influenza (n=5), CHIP (−) mild COVID-19 (n=5), CHIP (−) severe COVID-19 (n=3), and CHIP (+) severe COVID-19 (n=6) specimens (Supplementary Table 3 and 4, see Methods), with an average of 6,100 unique molecular identifies (UMIs), representing 1,400 genes. The reproducibility and quality were ensured (Extended Data Fig. 2a-c)^26^. Based on t-distributed stochastic neighbor embedding (t-SNE) of transcriptome profiles, 24 subgroups were derived. Assigning cell types with previously annotated marker genes (Extended Data Fig. 2d-e, see Methods)^26^, we focused on 9 major immune cell types, including IgG^+^ B cell, IgG^-^ B cell, CD4^+^ T cell, naïve CD4^+^ T cell, CD8^+^ T cell, CD8^+^ memory T cell, natural killer (NK) T cell, classical monocyte, and non-classical monocyte in subsequent analyses (Fig. 2a). Non-immune cells such as platelets, red blood cells (RBCs), and uncategorized small cell populations were excluded. Notably, the proportion of monocytes was markedly increased, particularly in CHIP (+) severe COVID-19 patients (K–S test, *P*=3.49e-3) (Extended Data Fig. 2f-g), consistent with the known impact of CHIP with increased myeloid cell population^27^.

**Figure 2.**
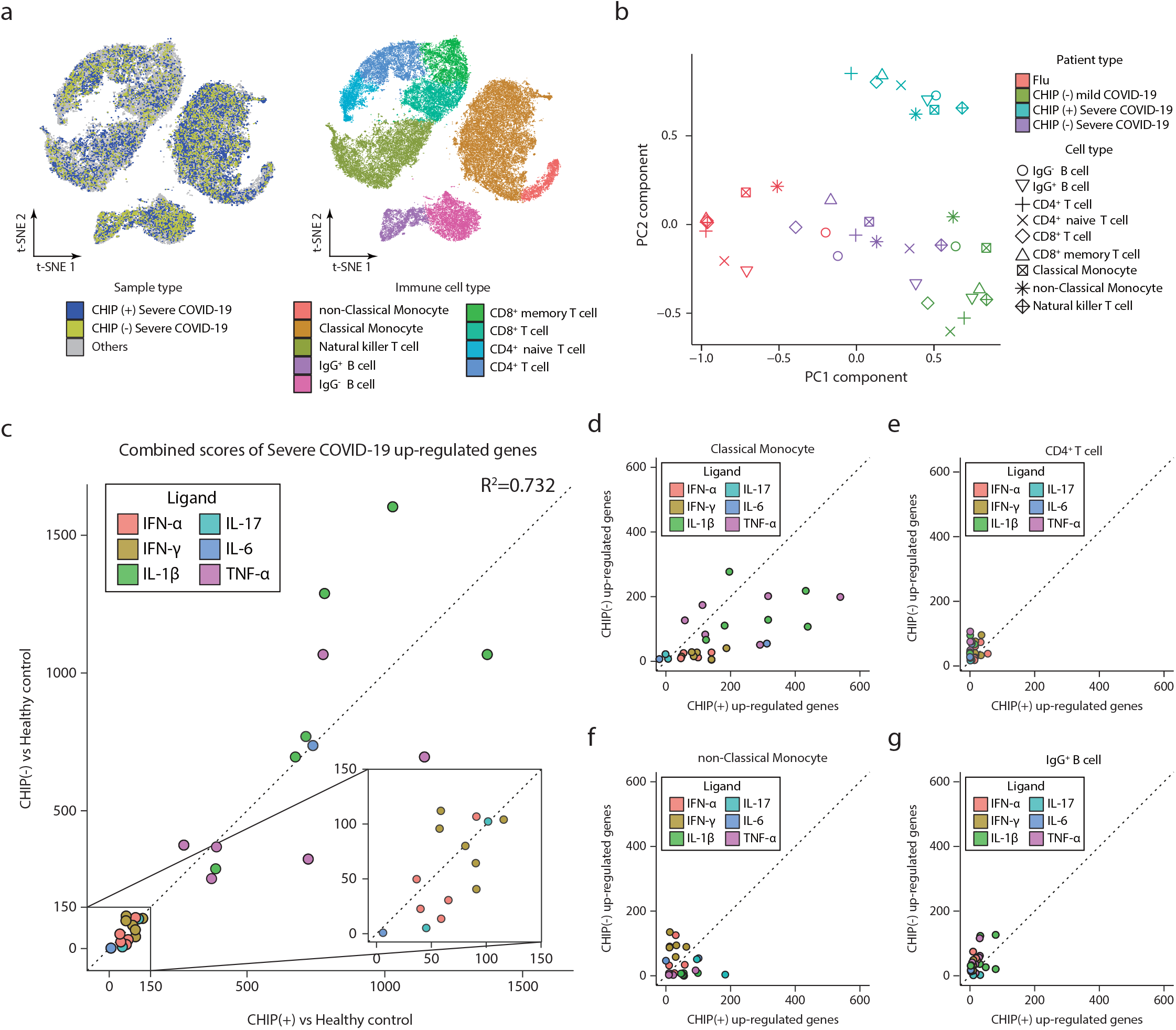
Single-cell transcriptome analyses of COVID-19 according to CHIP status. **a**, Scatter plots of integrated scRNA-seq data represented with a t-SNE method. Left, immunes cells are presented according to the presence of CHIP. Others group indicates CHIP (−) COVID-19 mild, influenza, and healthy normal control groups. Right, nine immune cell types were plotted by a t-SNE method. **b**, A PCA analysis with transcriptome profiles according to the immune cell types and disease groups. Colors and shapes represent the respective disease groups or cell types. **c**, A scatter plot showing combined scores of LINCS L1000 Ligand Perturbations up gene ontology library between CHIP (+) severe COVID-19 and CHIP (−) severe COVID-19 compared to the healthy normal control. Identity lines are presented on diagonal. Square of correlation of combined scores is shown together. **d-g**, Scatter plots showing combined scores same gene ontology library in Fig. 2c for differentially expressed genes between CHIP (+) and CHIP (−) severe COVID-19 in classical monocyte (d), CD4+ T cell (e), non-classical monocyte (f), and IgG + B cell (g). The horizontal axis, up-regulated genes in CHIP (+); Vertical axis, up-regulated genes in CHIP (−). Identity lines are presented on diagonal. The color indicates types of perturbed ligand.

In terms of transcriptome profiles at the cell-type resolution, as expected, most immune cell types originating from COVID-19 were clustered together when compared to influenza (Fig. 2b). Interestingly, those from severe COVID-19 were subdivided according to the presence of CHIP (Fig. 2b). To investigate relevant biological functions that establish such unique host immune responses in CHIP (+) severe COVID-19, we identified up-regulated genes specific to CHIP (+) severe COVID-19 compared to healthy normal control, influenza, CHIP (−) mild COVID-19, and CHIP (−) severe COVID-19, respectively, using MAST algorithm (Extended Data Fig. 3a)^28^. Regardless of both normal and disease control groups, tumor necrosis factor (TNF-α)/NF-kB and interferon-gamma (IFN-γ) responses were commonly enriched in up-regulated genes in CHIP (+) severe COVID-19 (Extended Data Fig. 3b). A noticeable enrichment of CHIP (+) up-regulated genes was observed in classical monocytes (Extended Data Fig. 3c).

To further investigate the unique immune signatures at the individual immune cell-type resolution, we conducted gene set enrichment analysis (GSEA) for differentially expressed genes between CHIP (+) and CHIP (−) severe COVID-19. Based on cytokine-responsive gene sets originated from each cytokine treated cells (LINC L1000 ligand perturbation analysis in Enrichr) (see Methods)^29^, strong IL-1β and TNF-α responses were observed in both CHIP (+) and CHIP (−) severe COVID-19 compared to healthy normal control (Fig. 2c). However, direct comparison between CHIP (+) and CHIP (−) severe COVID-19 revealed that immune responses in classical monocytes were strongly skewed towards CHIP (+) patients (Fig. 2d), while other cell types such as T cells, B cells or non-classical monocytes did not show such trend (Fig. 2e-g, Extended Data Fig. 4a-e). Taken together, CHIP (+) patients under COVID-19 infection present unique host immune responses, particularly in classical monocytes.

### Classical monocyte-mediated hyperinflammation in CHIP (+) severe COVID-19

To examine how the presence of CHIP attributes to the immunologic responses in classical monocytes, we focused on analyzing 445 up- and 417 down-regulated CHIP (+) specific genes. CHIP (+) up-regulated genes demonstrated enrichment with inflammation cytokine responses, such as IL-1β compared to CHIP (−) (Mann-Whitney’s U test, *P*=3.24e-2, IL-1β) (Fig. 3a). Other COVID-19 representative interleukins such as IL-6, IL-10 and IL-15 were also elevated in CHIP (+) severe COVID-19. (Fig. 3b)^30^. Notably, CHIP (+) up-regulated genes were strongly related to both type I and II interferon (IFN) responses (Mann-Whitney’s U test, *P*=1.08e-3, IFN-γ; *P*=3.97e-3, IFN-α). We previously proposed that a group of genes involved in type I IFN-induced TNF-α mediated hyperinflammation by abolishing the tolerance effects of TNF-α (Class 1 gene in Park *et al*.^31^) in monocyte has a critical role in promoting hyperinflammation of COVID-19^26^. Consistently, the CHIP (+) up-regulated genes presented a modest enrichment with the Class I genes (Fig. 3c), partly explaining the hyperinflammation signature of CHIP (+). However, other inflammatory TLR-induced genes regardless of TNF-α tolerization (Class II and III genes in Park *et al*.^31^) were extremely biased to CHIP (+) up-regulated genes, postulating the presence of additional classical monocyte driven responses exacerbating the inflammation signatures in CHIP (+).

**Figure 3.**
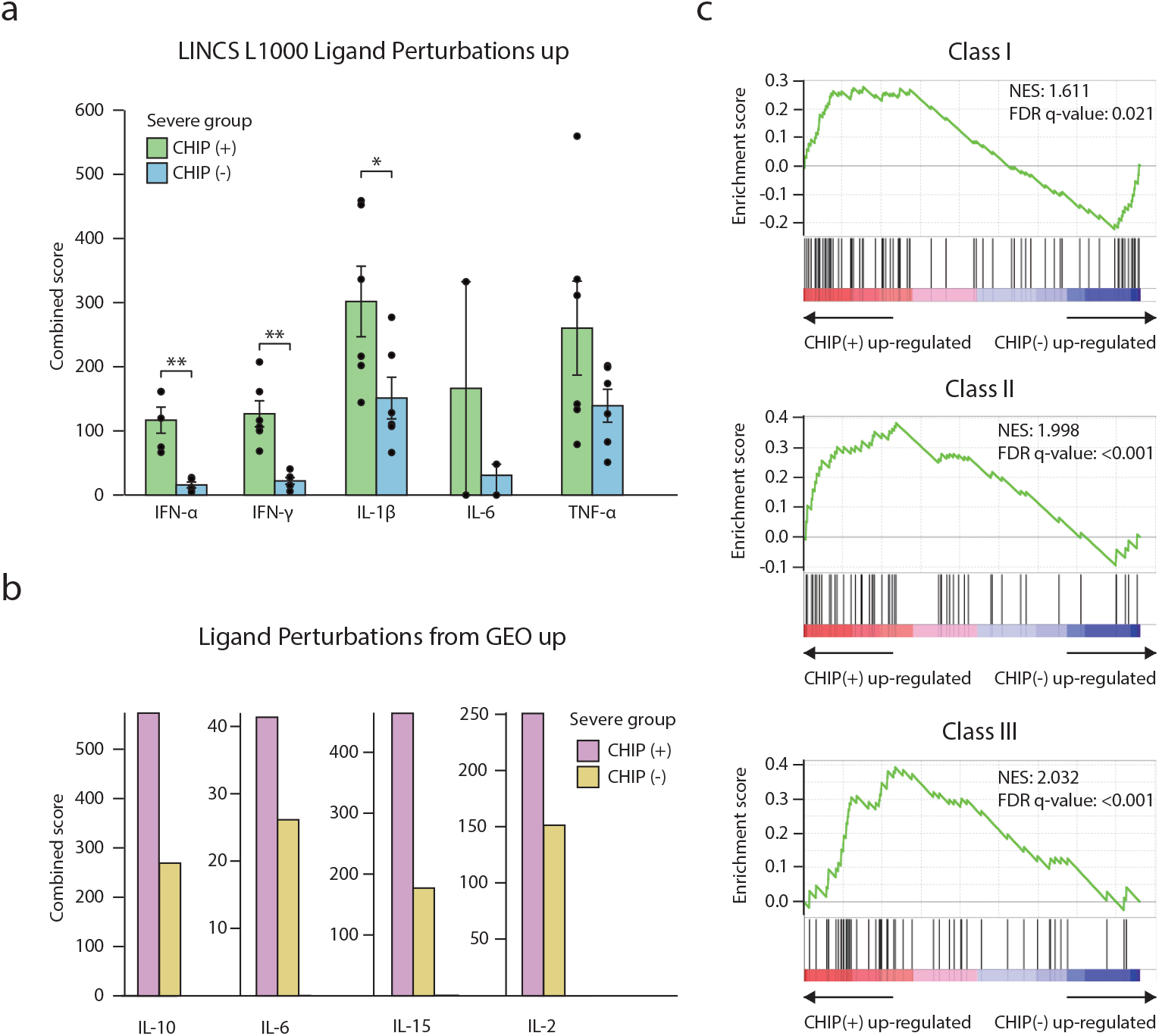
CHIP specific immune signatures in classical monocytes. **a**, Barplots showing combined scores of DEGs in classical monocytes between CHIP (+) and CHIP (−) severe COVID-19 for same gene ontology library in Fig. 2c-g. The color indicates up-regulated genes in CHIP (+) (green) and CHIP (−) (blue). Each point indicates one ligand perturbation term in the library, in total 6 different types of ligands: IFN-α (n=5), IL-1β (n=6), IFN-γ (n=6), IL-17 (n=2), IL-6 (n=2), TNF-α (n=6). The mean and standard error of the mean (s.e.m) of each gene set are shown together. One-sided Mann-Whitney’s U test was performed (*: *P<*0.05, **: *P<*0.01). **b**, Barplots showing combined scores of same gene sets in Fig. 3a for Ligand Perturbations from GEO up gene ontology library. The color indicates up-regulated genes of CHIP (+) (pink) and CHIP (−) (yellow). **c**, Gene set enrichment analysis (GSEA) on the same DEGs used in Fig. 3a-b for three classes of TLR induced genes^31^. Each plot shows the distribution of enrichment of distinct class genes along with the list of the DEGs pre-ranked with log-fold changes based on the CHIP (−) severe COVID-19. The color under the plot indicates patient groups of up-regulated genes. Red, up-regulated in CHIP (+) severe COVID-19. Blue, up-regulated in CHIP (−) severe COVID-19. Normalized enrichment scores (NES) and FDR are shown together. Top, Class I genes. Middle, Class II genes. Bottom, Class III genes.

### IFN-γ mediated hyperinflammation in CHIP (+) severe COVID-19 revealed by a pseudotime analysis

Intriguingly, IFN-γ response was notably high in CHIP (+) up-regulated genes (Fig. 3a). As a high level of IFN-γ has been reported as an indicator of severe COVID-19^8,32,33^, we hypothesized that IFN-γ response could attribute to hyperinflammation signatures of classical monocytes in CHIP (+) patients. We first determined the association between IFN-γ response and COVID-19 severity in CHIP (+) by conducting pseudotime analysis using the specimens collected twice from one patient to exclude innate individual biases (Fig. 4a) (see Methods). After ordering cells along with the trajectory analysis, we allocated the annotation of high and low inflammation clusters based on inflammatory signatures termed in MsigDB Hallmark 2020 (Fig. 4b). We found that IFN-γ response genes were significantly enriched in a high inflammation group of CHIP (+), even more potent than that of CHIP (−) (Fig. 4c).

**Figure 4.**
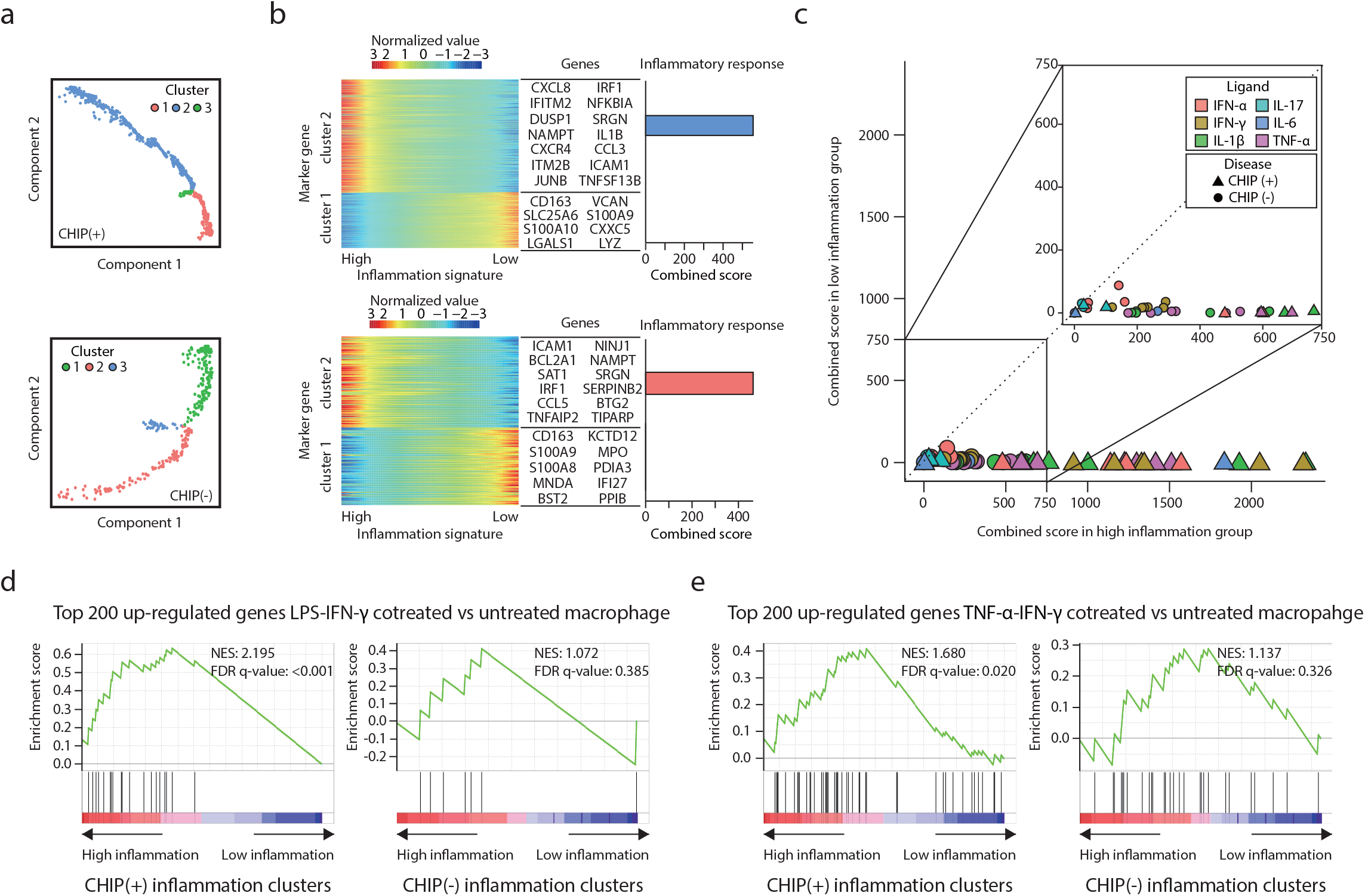
Pseudotime analyses of severe COVID-19 with CHIP in classical monocytes. **a-c**, Pseudotime analyses for CHIP (+) or CHIP (−) severe COVID-19 patients. **a**, Cells are aligned according to the pseudotime axis calculated by Monocle2. The color indicates a type of cluster. Top, CHIP (+) patient. Bottom, CHIP (−) patient. **b**, Heatmaps representing the expression level of marker genes for the ordered cells. The list of genes is the representative gene set of each cluster. Barplots showing combined scores of each cluster for inflammatory response term in MsigDB Hallmark 2020. **c**, Scatter plots of combined scores of marker genes of inflammation clusters for LINCS L1000 Ligand Perturbations up gene ontology library. The horizontal axis, high inflammation cluster; the vertical axis, low inflammation cluster. Identity lines are presented on diagonal. Colors indicate types of perturbed ligand for IFN-α (n=5), IL-1β (n=6), IFN-γ (n=6), IL-17 (n=2), IL-6 (n=2), TNF-α (n=6). Shapes represent the presence or absence of CHIP. **d, e**, GSEA plots for marker genes of inflammation clusters in CHIP (+) and CHIP (−), respectively. Genes are ordered based on log-fold changes between high inflammation cluster and low inflammation cluster. Normalized enrichment scores (NES) and FDR are presented for DEGs between M1-like (LPS-IFN-γ stimulated) and M0 macrophage (untreated) ^36^(d) and DEGs between TNF-α-IFN-γ co-treatment and untreated condition^35^ (e), respectively. LPS indicates Lipopolysaccharide.

Recent mouse and scRNA-seq comparison studies have highlighted the immuno-pathogenic contribution of IFN-γ in severe COVID-19 as demonstrated by IFN-γ induced inflammatory macrophage phenotype and synergistic effect of IFN-γ and TNF-α^34,35^. Consistently, in our analysis, CHIP (+) specific genes were significantly enriched by pro-inflammatory M1-like macrophage-specific genes obtained from two independent studies^36 37^(Fig. 4d and Extended Data Fig. 5a) and associated with up-regulated genes by co-treatment of TNF-α and IFN-γ (Fig. 4e)^35^. Such enrichment suggests that CHIP (+) severe COVID-19 patients are representative cases that could be explained by previously discovered IFN-γ mediated disease exacerbating mechanism. For a treatment perspective, we noticed that spleen tyrosine kinase (Syk) inhibitor, which is known to reduce the expressions of interferon-stimulated genes^38^, may be an effective molecule for the intervention of CHIP (+) up-regulated genes (Paired t-test, *P*=0.042) (Extended Data Fig. 5b). Taken together, IFN-γ response is thought to be a main source of the hyperinflammation immune signature in CHIP (+) severe COVID-19.

### *DNMT3A* mutation-specific hypo-DMRs are linked to IFN-γ response genes

Next, we sought to identify mechanisms by which IFN-γ response genes are up-regulated explicitly in CHIP (+) COVID-19 patients. Considering mutations of multiple epigenetic regulators such as *DNMT3A, TET2*, and *ASXL1* in CHIP, we hypothesized that altered chromatin activity of *cis*-regulatory elements might exert CHIP-specific gene expression. To test our hypothesis, we identified 2,348 differentially methylated regions (DMRs) in acute myeloid leukemia (AML) patients carrying *DNMT3A* mutations (see Methods) (Supplementary Table 5)^39^. In support of our hypothesis, these DMRs were highly overlapped with putative regulatory elements (Fisher’s exact test, *P*<0.001, Proximal to the promoter; *P*<0.001, Distal regulatory element) (Fig. 5a) (see Methods). As exemplified in *FOXO3* and *NFIL3*, known IFN response genes, CHIP (+) up-regulated genes were also more closely located to the hypo-DMRs (Fig. 5b, Extended Data fig. 6).

**Figure 5.**
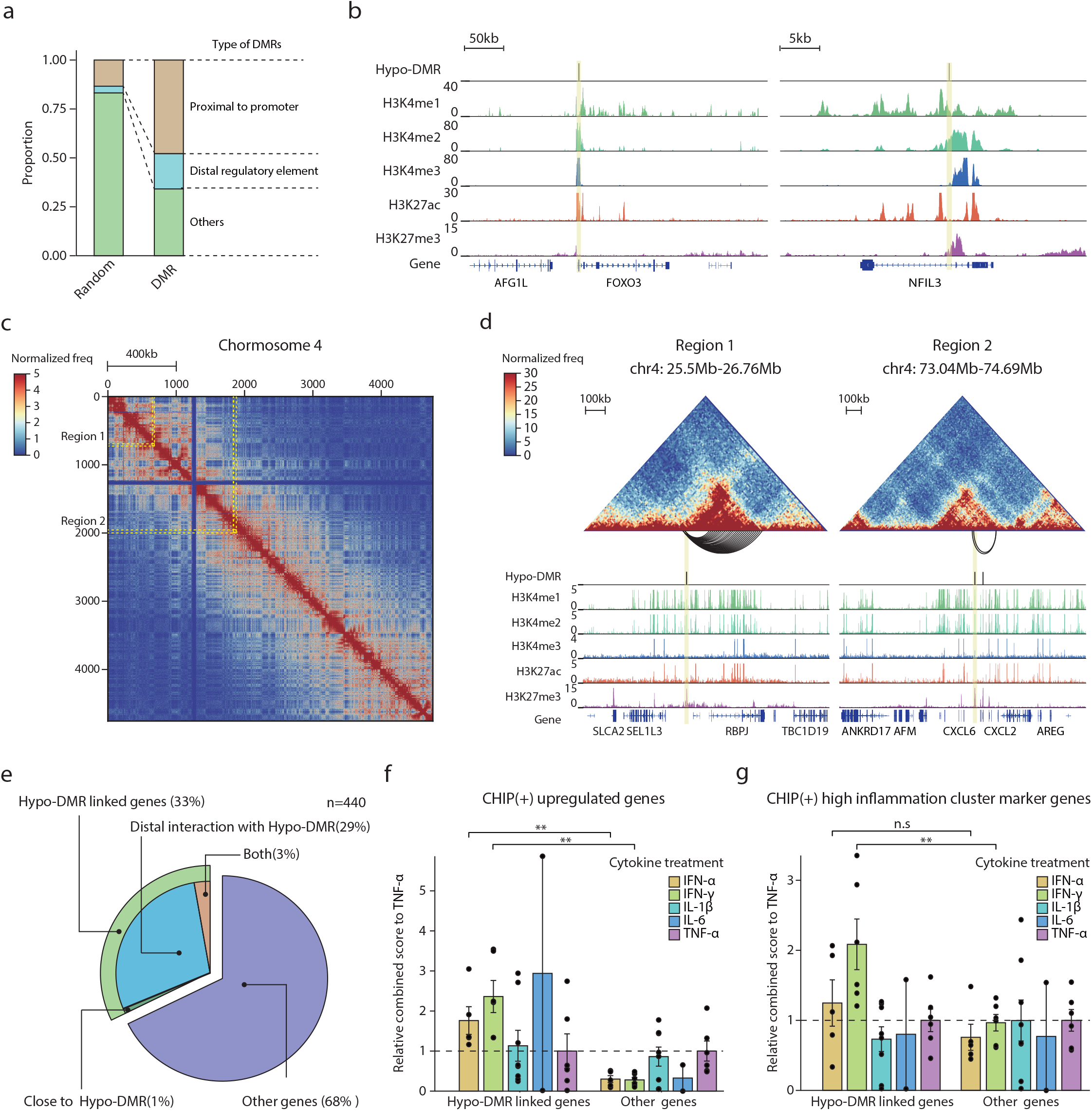
Regulatory potential of CHIP-specific hypo-DMRs in IFN response genes. **a**, Stacked bar plots showing the proportion of annotated hypo- and hyper-DMRs. In DMR, Proximal to the promoter, n=1124; Distal regulatory element, n=423; others, n=801. In random for DMR, Proximal to promoter, n=314; Distal regulatory element, n=81; others, n=1953. **b**, Examples of CHIP (+) up-regulated genes are presented with hypo-DMRs and multiple histone modification signatures for *FOXO3* (right) and *NFIL3* (left). **c**, Examples of analysis of Hi-C data. Whole interaction map of chromosome 4 in 40kb resolution. The color indicates normalized chromatin contact frequencies. Two example regions containing CHIP (+) severe COVID-19 up-regulated genes in classical monocytes (region 1 and region 2) are highlighted by yellow dashed lines. **d**, Hi-C contact maps (heatmap) for region 1 (*RBPJ*) and region 2 (*CXCL2*) are shown together with significant long-range chromatin interactions with the promoter regions (arcs on the below of the heatmap) and distribution of hypo-DMRs and histone modifications signals. **e**, A pie chart showing proportions of hypo-DMR linked up-regulated genes in CHIP (+) severe COVID-19 classical monocytes. The color indicates the types of linkage between hypo-DMRs and up-regulated genes. **f, g**, Barplots of relative combined scores to TNF-α of genes linked to hypo-DMRs and others for same gene ontology library in Fig. 2c-g. Colors indicate types of perturbed ligands. Each point indicates one ligand perturbation term. The mean and standard error of the mean (s.e.m) of each gene set are presented in barplot. One-sided Mann-Whitney’s U test was performed (n.s: non-significant, **: *P<*0.01). Up-regulated genes in CHIP (+) compared to CHIP (−) (f) and marker gens of high inflammation cluster compared to low inflammation cluster of CHIP (+) patient in Fig. 4 (g).

As many *cis*-regulatory elements are known to target genes over large genomic distances^40^, we performed *in situ* Hi-C experiments on CD14^++^/CD16^-^ classical monocytes of two healthy donors to precisely annotate target genes of CHIP-dependent DMRs (see Methods). With ∼500M long-range chromatin interactions over 15kb genomic distance, we defined significant long-range chromatin contacts using covNorm in 10kb resolution (see Methods)^41^. Using this information, as illustrated in *RBPJ* and *CXCL2* genes (Fig. 5c-d), we revealed that, in total, around 33% of CHIP-specific up-regulated genes, denoted as ‘linked genes’, were associated with hypo-DMRs either in proximal (within 15kb) or long-range chromatin interactions (over 15kb but less than 2Mb) (Fig. 5e). Notably, those linked genes were more enriched by both type I and II IFN response genes compared to the remaining up-regulated genes (Mann-Whitney’s U test, *P*=2.50e-3, IFN-γ; *P* =3.97e-3, IFN-α) (see Methods) (Fig. 5f). Regarding high inflammation cluster in pseudotime analysis, we revealed that IFN-γ response genes largely overlap by hypo-DMRs linked genes (Mann-Whitney’s U test, *P* =1.08e-3, IFN-γ) (Fig. 5g). Our results strongly support that CHIP-dependent altered chromatin activities are associated with putative *cis*-regulatory elements of IFN-γ response genes, which may establish unique gene expression profiles in CHIP patients during immune responses.

### Activation of poised and bivalent *cis*-regulatory elements primes IFN-γ response genes

To further characterize CHIP-specific hypo-DMRs, we examined the enrichment of four representative histone modification marks including histone H3 4^th^ lysine mono-methylation (H3K4me1), tri-methylation (H3K4me3), 27^th^ lysine acetylation (H3K27ac), and tri-methylation (H3K27me3) of primary human classical monocyte^42,43^. When comparing DMRs to randomly selected genomic regions, hypo-DMRs were mostly marked by H3K27me3 in the primary human classical monocytes, while hyper-DMRs were enriched by H3K27ac as an indicator of active regulatory elements (Fig. 6a-b). However, interestingly, a subset of hypo-DMRs, linking to CHIP (+) up-regulated genes, were also significantly co-occupied by H3K4me1 and H3K4me3 peaks compared to the unlinked hypo-DMRs (Fisher’s exact test, *P*<0.001, H3K4me1; *P*<0.001, H3K4me3; *P* =2.37e-2, H3K27ac) (Fig. 6c-d). Such co-exhibition of inactive and active chromatin signatures indicates that the regulatory elements of CHIP (+) up-regulated genes have shifted from poised or bivalent status to an active chromatin state through the process of CHIP dependent hypo-DNA methylation. To test this possibility, we annotated chromatin states of linked hypo-DMRs according to the combination of the histone modification marks (see Methods). We found that promoter-distal hypo-DMRs were significantly enriched by poised enhancer chromatin signatures (H3K4me1 without H3K27me3) (Fisher’s exact test, *P* <0.001) (Fig. 6e). Similarly, promoter-proximal hypo-DMRs were enriched by bivalent promoters (H3K4me3 and H3K27me3) or active promoters (H3K4me3) (Fisher’s exact test, *P* = 3.80e-3, Active promoter; *P*= 7.36e-2, Bivalent) (Fig. 6f). Thus, CHIP mutants appear to reprogram the epigenetic states including the loss of silent marker at poised enhancers or bivalent promoters, which prime the IFN associated immune response genes, thereby driving hyperinflammation and leading to the critical course of COVID-19 (Fig. 7).

**Figure 6.**
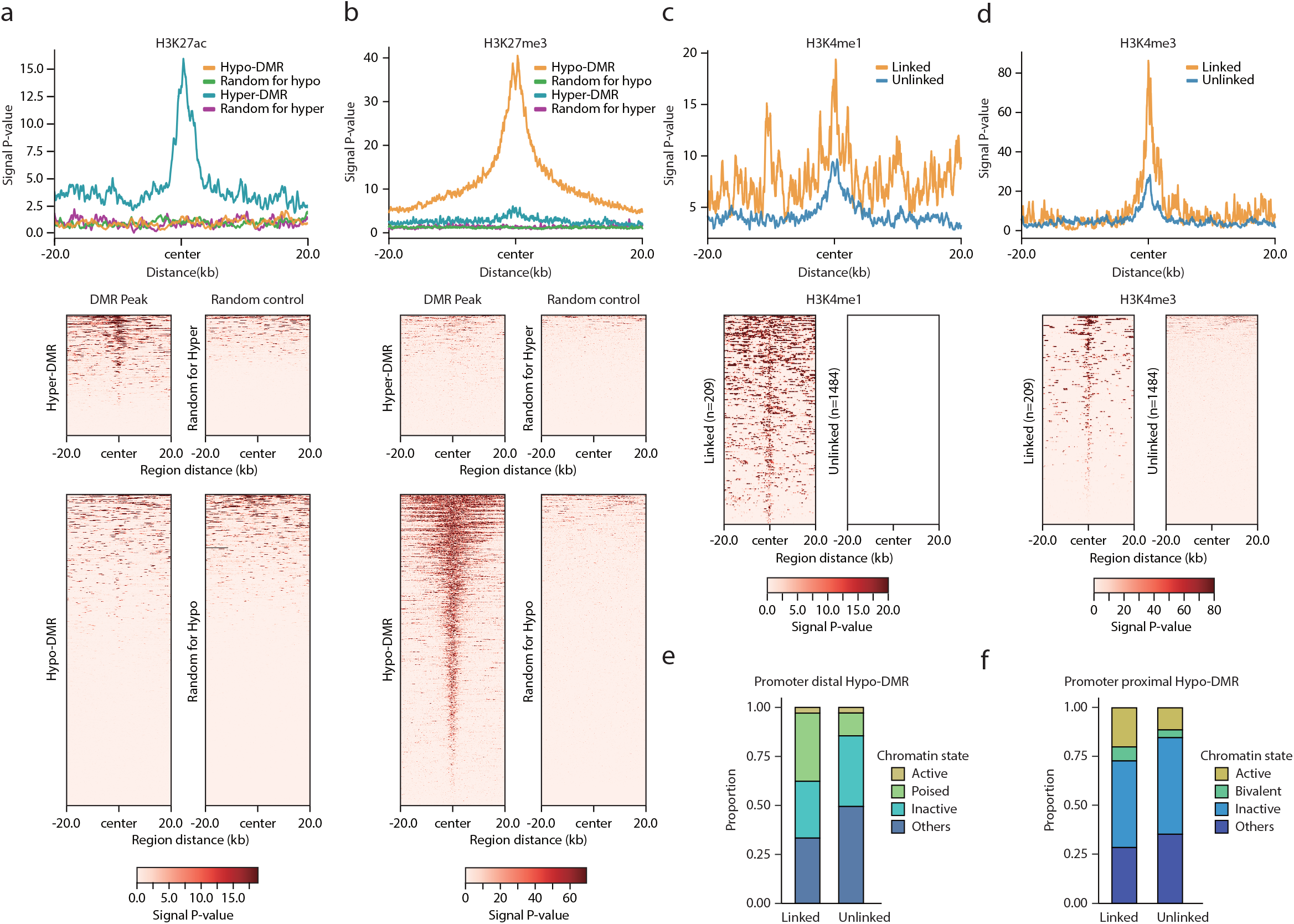
Chromatic states of hypo-DMRs linking to CHIP specifically up-regulated genes. **a, b**, CHIP-seq signal distribution (a for H3K27ac and b for H3K27me3) of 20k upstream and downstream surrounding of hypo-DMRs (n=1,693), hyper-DMRs (n=655) and randomly selected regions. Each randomly selected regions have the same distribution of chromosome number and length of hypo- and hyper-DMR, respectively. Top, average ChIP-seq signal distributions. Bottom, heatmaps of CHIP-seq signals of the corresponding regions. **c, d**, CHIP-seq signal distribution for CHIP (+) up-regulated genes linked (n=209) and unliked (n=1484) hypo-DMRs for H3K4me1 (c) and H3K4me3 (d). Top, average profiles of CHIP-seq signals for linked and unlinked hypo-DMRs. Bottom, heatmaps of CHIP-seq signals of the corresponding regions. **e, f**, Stacked barplots of the linked and unlinked hypo-DMRs with annotation of chromatin states. For statistical significance, one-sided Fisher’s exact test was performed between linked and unlinked for promoter-distal (e) or -proximal (f) hypo-DMRs.

**Figure 7.**
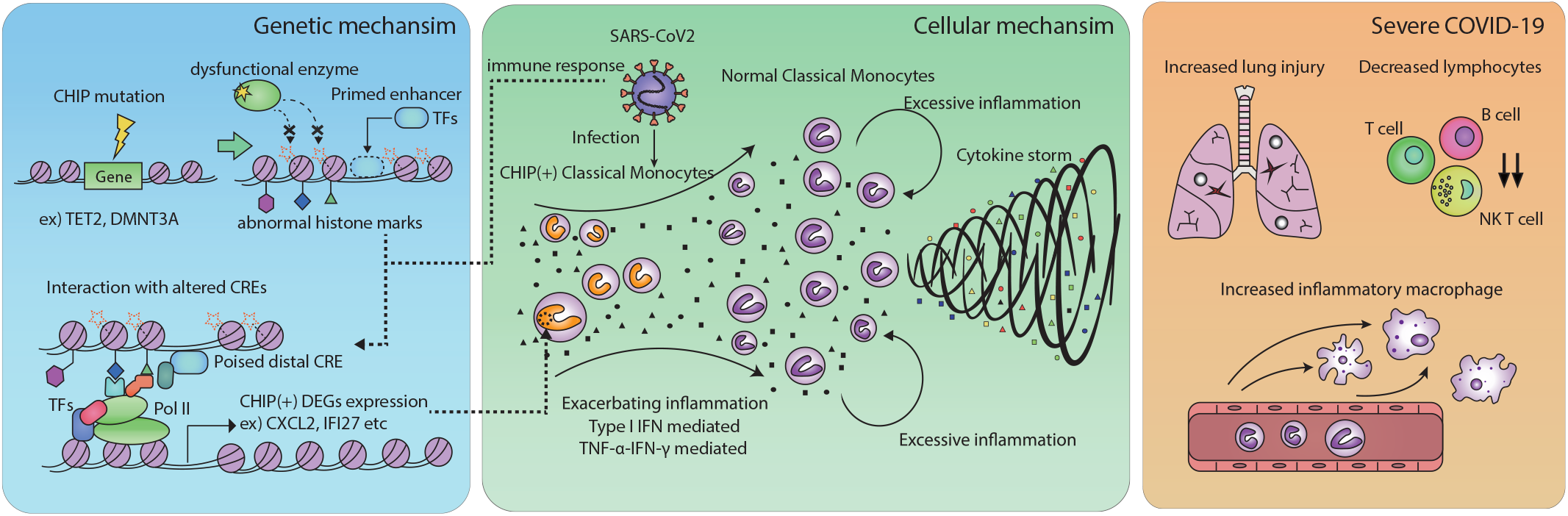
A proposed model for the pathogenesis of severe COVID-19 in patients with CHIP. A schematic overview shows the progression of disease exacerbation in terms of gene regulation, cellular level, and clinical/immunological signatures under CHIP (+) severe COVID-19.

## Discussion

By analyzing thorough clinical, radiological, and laboratory characteristics, as well as the presence of CHIP, we showed that CHIP is a novel risk factor for severe COVID-19 in previously healthy population without canonical clinical risk factors. scRNA-seq analysis revealed a distinct IFN-γ mediated immuno-pathogenic mechanism in CHIP (+) severe COVID-19 plausibly attributed by CHIP dependent chromatin reorganization. These results consistently indicate that CHIP may play a critical role in the progression of severe COVID-19, especially in previously healthy patients with its own immunologic pathway.

Exploration of an additional risk factor for severe COVID-19 is clinically valuable in this pandemic to predict the progression of COVID-19 more accurately and to improve our management strategy. Owing to its pro-inflammatory nature, CHIP may contribute to the progression of severe COVID-19. By analyzing more than 500 patients, Bolton *et al*. reported that CHIP is significantly associated with severe COVID-19^23^, especially for patients carrying non-putative driver mutations. There have been a few controversial reports, however, they needed to be critically appraised because of methodological considerations. Duployez *et al*. showed a higher prevalence of CHIP in severe COVID-19 patients than age-matched hematologic malignancy-free cohort, but they could not show that CHIP affects clinical outcome^24^. However, they could only analyze severe patients whose clinical outcomes might hardly be differentiated by solely the presence of CHIP. In another study conducted by Hameister *et al*. involving 102 hospitalized patients with COVID-19, the presence of CHIP was not associated with severe COVID-19^25^. But the study lacked a stratified analysis or an adjustment for possible interactions. By introducing a hierarchical clustering, followed by thorough statistical examinations including interaction analysis, we could clearly show that CHIP is an independent risk factor for severe COVID-19.

COVID-19 patients were reported to show heterogeneous symptoms ranging from asymptomatic to critical illness^3,44^. In line with it, many studies divided COVID-19 patients into subgroups defined by immunological characteristics, for instance, patterns of sepsis^45^, subpopulations of lymphocytes^44^, IFN responses in lung^46^, or carrying loss-of-function variants^47^. Single-cell techniques have been vigorously applied in COVID-19 to dissect underlying causes of the diverse immune responses^33,48,49^ and to elaborately determine the relationship between immune subtypes and clinical characteristics^50,51^. Despite those important works, none of the single-cell study has characterized the immunological effects of CHIP in COVID-19 yet. In this regard, the current study uniquely demonstrated how CHIP-associated somatic mutations in immune cells could actively establish a novel subgroup in COVID-19 patients. With single-cell immune transcriptome analysis, we could reveal that IFN-γ related hyperinflammation is a hallmark of CHIP (+) severe COVID-19. Especially, there was an enrichment of inflammatory signature in classical monocytes, which is compatible with recent knowledge regarding the effect of CHIP on myeloid askew hematopoietic stem cell differentiation^27^. From a cytokine perspective, IFN-γ and its synergism with TNF-α were thought to play a critical role in the pathogenesis of severe COVID-19 in CHIP (+) patients. This finding aligns with the previous report stating the role of IFN-γ and/or TNF-α in exacerbating chronic inflammatory disease by CHIP^52^. Our study also implies that not only type I IFN response but also type II IFN response plays an important role in disease exacerbation in certain patients with severe COVID-19. Additional interesting immunological finding reasonably explained with the CHIP biology is the up-regulation of genes related to inflammatory macrophage in CHIP (+) severe COVID-19. CHIP is well known to drive hyperinflammation in chronic disease mainly attributable to the altered function of monocyte and macrophage ^27^.

Recent studies have revealed the pathological effects of CHIP such as atherosclerosis and malignancy^18,53,54^ and characterized the effect of CHIP mutations on hematopoiesis and immunological functions regarding epigenetic mechanisms^55,56^. However, there is still an ambiguity in how physiological pathogenic characteristics are linked to the altered chromatin activities by CHIP mutations in actual patient cohorts. In our study, such linkage is explained as regulatory interactions between pathogenic genes and their *cis*-regulatory elements with the altered chromatin states, which are expected to be mediated by mutations in chromatin regulators (Fig. 7). Although further investigation of other types of CHIP mutations such as *TET2* and *ASXL1* are required to comprehensively characterize CHIP mutation-dependent epigenetic reprogramming, the pathogenesis of other CHIP-dependent diseases could be elucidated by considering the similar mechanisms.

There are several limitations in the present study. First, sample sizes of either entire cohort or scRNA-seq-analyzed patients were limited. Second, the proportion of severe COVID-19 was high in the present study since it was conducted mainly in tertiary-care hospitals. Although we could show an apparent incline to severe COVID-19 in CHIP (+) patients and its distinctive immune signature in this study, further validations in a larger prospective cohort representing general COVID-19 patients are warranted. Third, scRNA-seq was performed with PBMCs instead of lung fluid or infected lung tissues limiting our analysis in exploring systemic inflammation as a result of COVID-19 pulmonary disease. Lastly, further studies are needed to reveal the exact epigenetic differences between mutant and wild type monocytes in CHIP patients and the real time point when CHIP mutant monocytes stir hyperinflammation.

Lastly, in terms of therapeutic strategy, Syk inhibitors have been implicated in suppressing these pathogenic immune responses (Extended Data Fig. 5b) induced by CHIP. An *in vitro* study on Syk inhibitor fostamatinib suggested its therapeutic effect against COVID-19^57^, and we have pending results of a randomized placebo-controlled trial with the drug (NCT04579393). As the therapeutic efficacy of Syk inhibitors could be more potent in CHIP (+) patients, it is worth evaluating treatment outcomes involving Syk inhibitors according to CHIP status.

In summary, despite such limitations, we successfully clarified that there is a distinct CHIP-driven severe COVID-19 subgroup and elucidated its unique immunological mechanism. Revealing the underlying epigenetic mechanism for the altered immune function that aligns with well-known CHIP biology suggests the robustness of our findings. It appears that classical monocytes in patients with CHIP (+) COVID-19 undergo distinct immune responses, thus focusing on immunomodulation strategies according to the presence of CHIP is required. Considering the shared pathogenic host immune response across infections, we postulate that our findings might bring a better understanding of previously unexplained exacerbation of clinical conditions by various viruses.

## Acknowledgments

We thank all our laboratory members for their support and critical suggestions throughout this work. Funding: This work was funded by the SUHF Fellowship (to I.J.). We also thank Dr. Shin for sharing specimens to determine the presence of CHIP.

## Author Contribution

CKK, BC, YK, and IJ conceived the study. SP generated scRNA-seq data. AJL and YK generated Hi-C data. CKK, BC, SK, SHY, DL, and SK performed data analysis. CKK, EC, JJ, PGC, WBP, ESK, HBK, NJK, MO, SK, HI, JK, YHL, JL, JYL, JHM, and K-HS contributed to the collection of clinical information and samples. CKK, BC, YK, and IJ contributed to data interpretation with assistant from SK and KK. CKK and BC prepared the manuscript with assistance from YK and IJ. All authors read and commented on the manuscript.

## Figure Legends

**Extended Data Figure 1.**
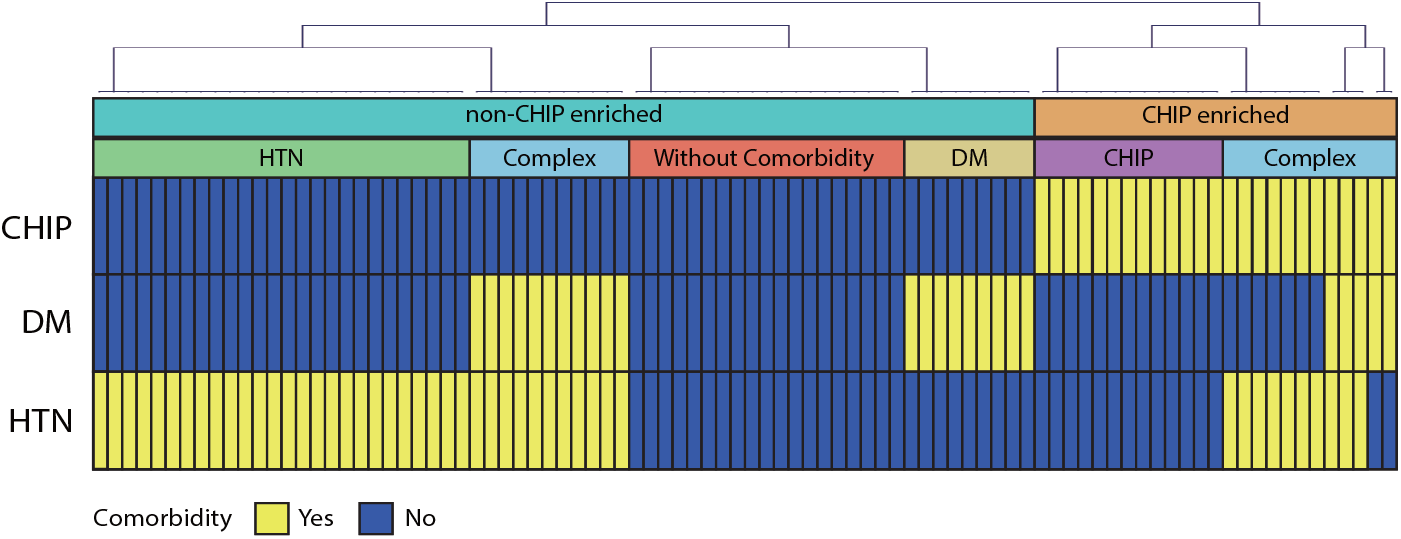
A heatmap showing hierarchical clustering of severe COVID-19 patients with comorbidities and CHIP. The first bar represents two clusters, cyan for non-CHIP enriched (n=65), and orange for CHIP enriched one (n=25). The second bar represents five distinct types of sub-clusters, namely, red, for without comorbidity (n=19), orange for DM (n=9), green for HTN (n=26), purple for CHIP (n=13), and blue for complex (n=23). The color in the heatmap represents the presence of CHIP.

**Extended Data Figure 2.**
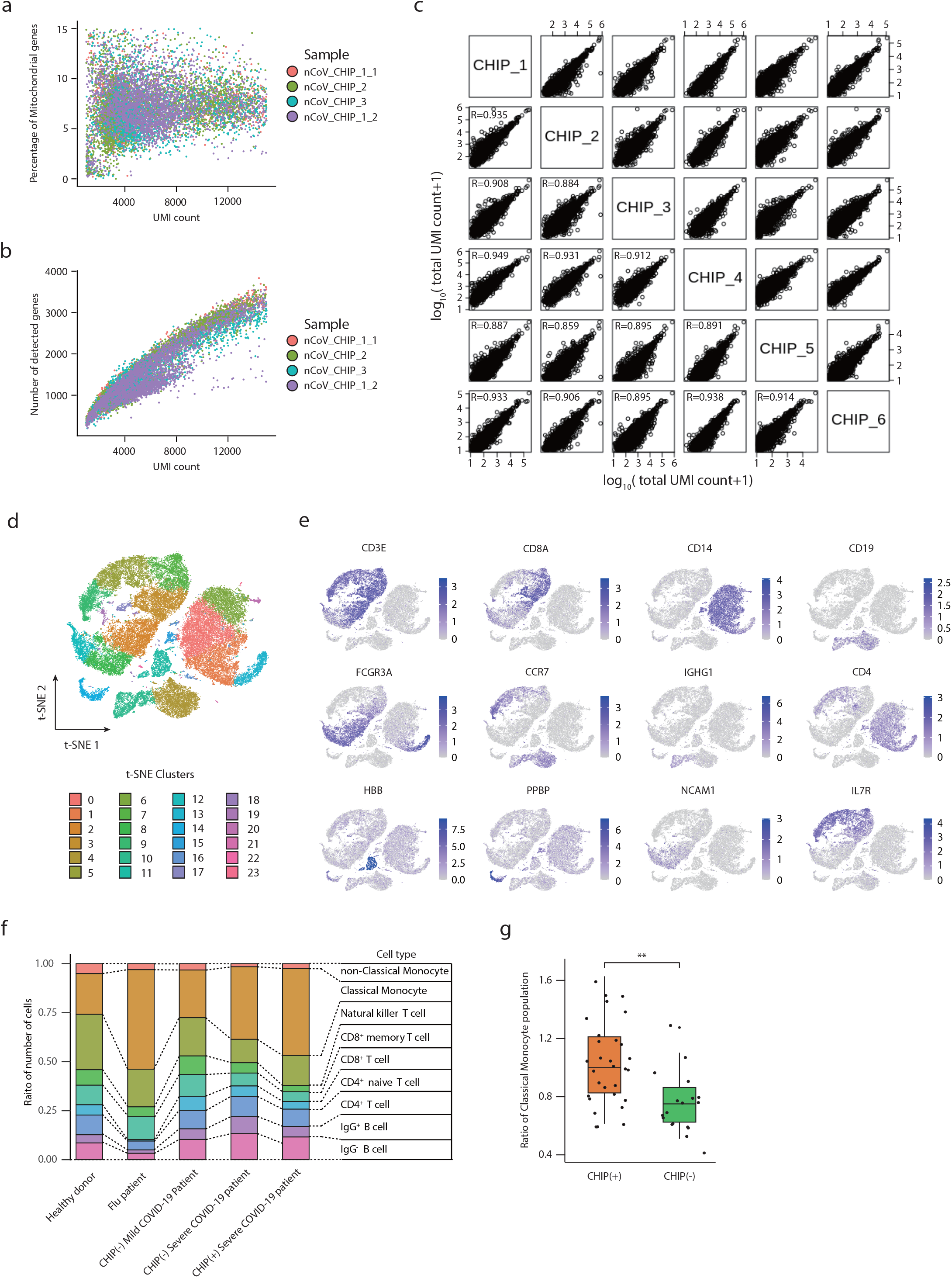
Quality-control and cell-type annotation of single-cell RNA-seq results. **a, b**, Scatter plots showing CHIP (+) patients’ immune cells according to the UMI count and other features. Percentage of the mitochondrial genes (a) and the number of detected genes (b). **c**, Scatter plot of log 10-transformed count data between individual patients. R represents Pearson’s correlation coefficient values. **d**, t-SNE clusters of integrated single-cell data. **e**, Each cell type marker gene expression on t-SNE plot. **f**, Stacked barplots show cell-type proportions of each patient group. **g**, Boxplots showing the ratio of the proportion of classical monocytes. (left) All pairs within CHIP (+) patients (n=30). (right) all pairs between CHIP (+) and CHIP (−) (n=18). The box represents the interquartile range (IQR) and the whiskers correspond to the highest and lowest points within 1.5 × IQR. Two-sided Kolmogorov-Smirnov test were performed (** < p-value 0.01).

**Extended Data Figure 3.**
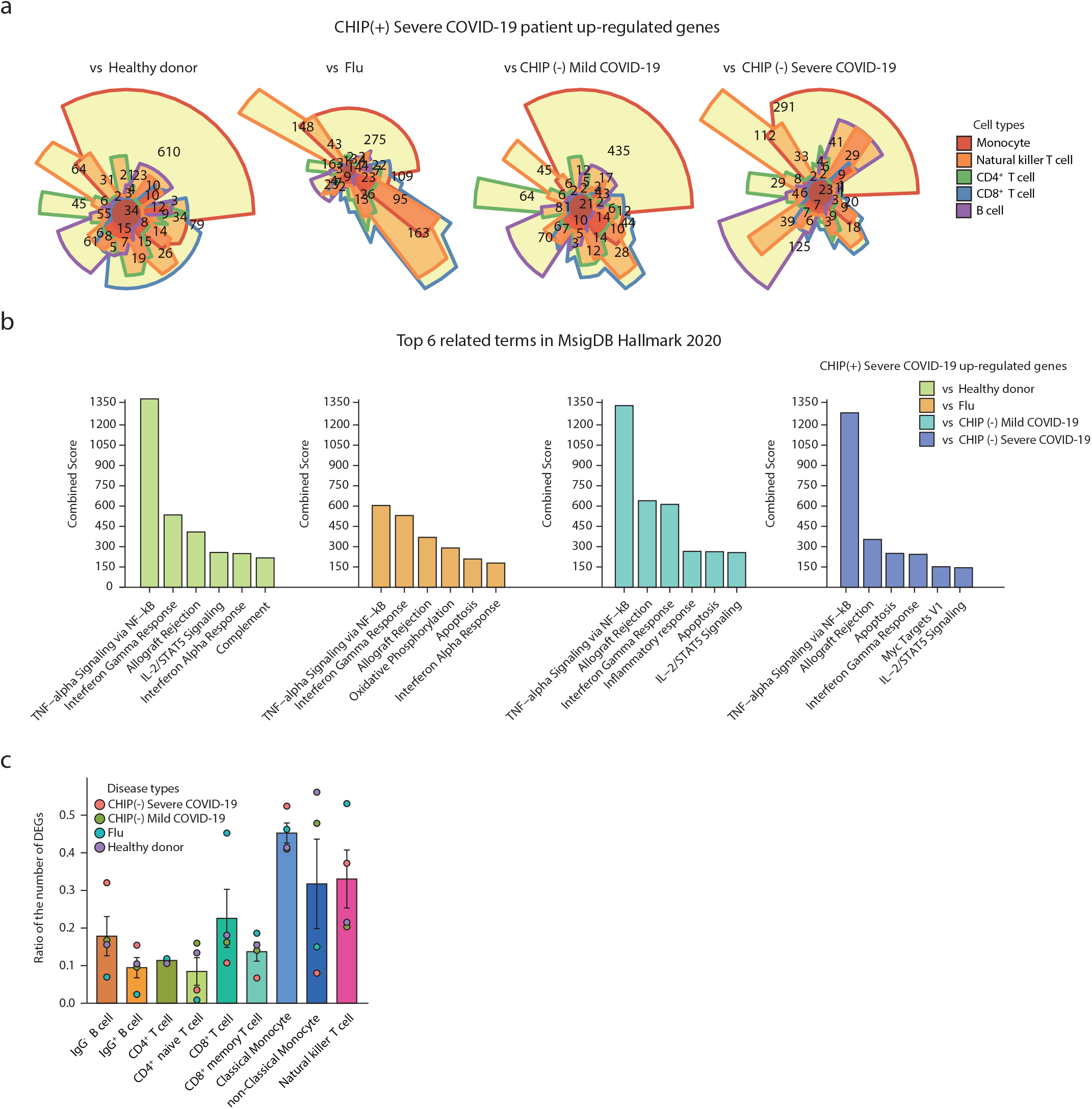
Analyses of differently expressed genes between CHIP (+) severe COVID-19 and other groups. **a**, Chow-ruskey venn diagram of the up-regulated genes in CHIP (+) severe COVID-19 compared to other disease groups. The color of the border in diagrams represents five immune cell types. Monocyte (red), classical monocyte and non-classical monocyte; natural killer T cells (orange); CD4^+^ T cell (green), CD4^+^ T cell and CD4^+^ naïve T cell; CD8^+^ T cell (blue), CD8^+^ T cell and CD8^+^ memory T cells; B cell (purple), IgG^+^ B cell and IgG^-^ B cell. **b**, Barplots showing combined scores for gene ontology terms in MsigDB Hallmark 2020 with up-regulated genes in CHIP(+) severe COVID-19. Top six terms were presented. The color indicates normal or disease controls. **c**, Ratio of cell-type specific CHIP (+) severe COVID-19 up-regulated genes compared to total up-regulated genes for comparison with other disease groups. The mean and standard error of the mean (s.e.m) of each gene set are presented in barplots. The color indicates the type of disease group for comparison.

**Extended Data Figure 4.**
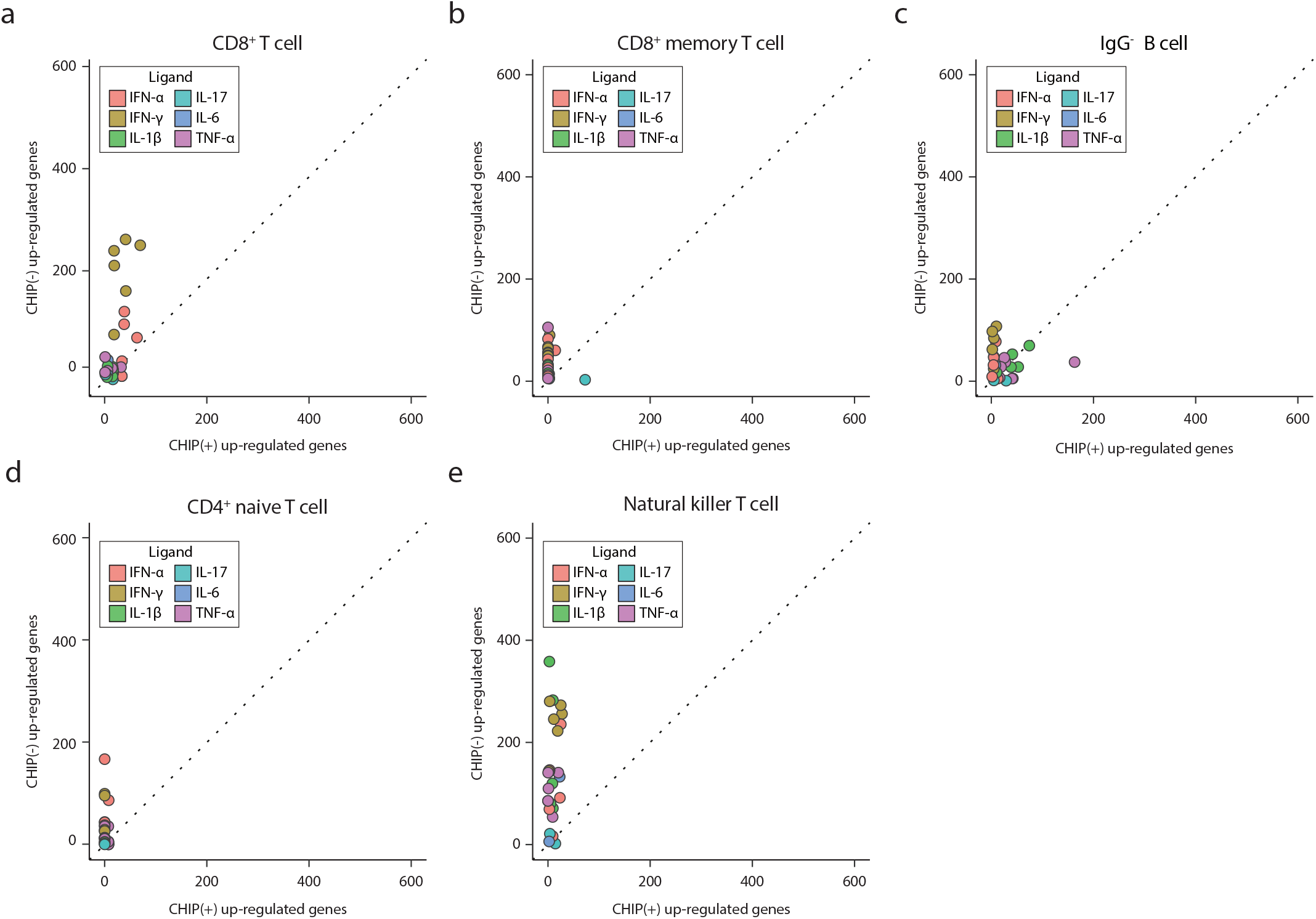
Enriched immune signatures in CHIP (+) and CHIP (−) severe COVID-19 up-regulated genes compared to healthy normal control. **a-e**, Scatter plots showing combined scores of differentially expressed genes (DEGs) in selected cell types between severe COVID-19 patient groups for same gene ontology library in Fig. 2c. The horizontal axis, up-regulated genes in CHIP (+) patients; the vertical axis, up-regulated genes in CHIP (−) patients. Identity lines are presented on diagonal. The color indicates types of perturbed ligand in database terms. **a**, CD8+ T cell. **b**, CD8+ memory T cell. **c**, IgG-B cell. **d**, CD4+ naive T cell. **e**, Natural killer T cell.

**Extended Data Figure 5.**
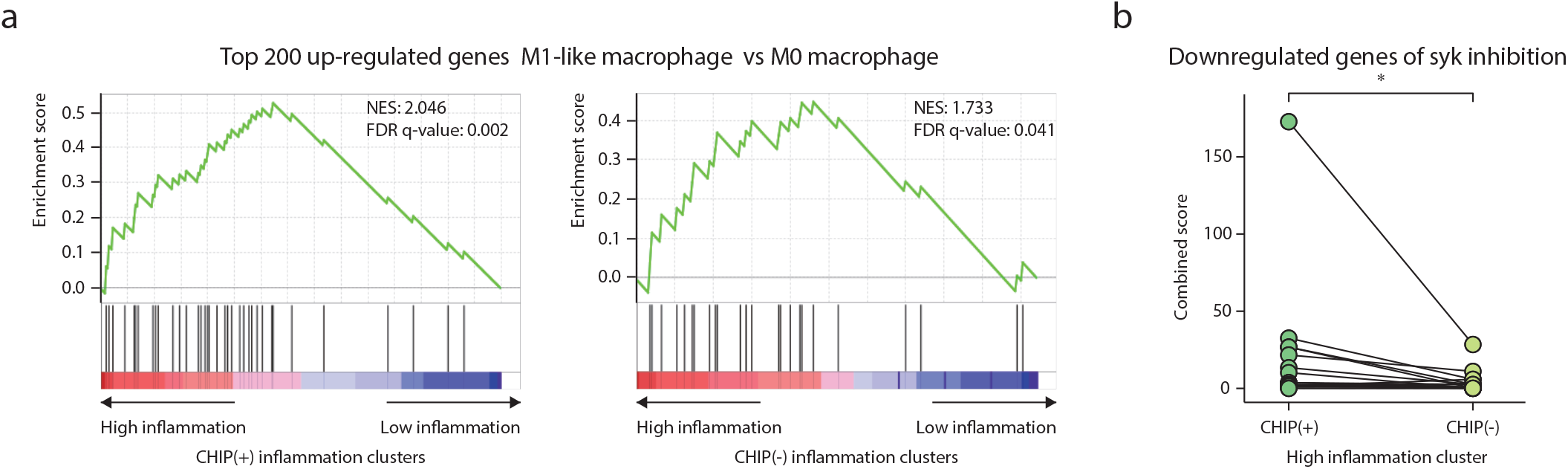
An inflammatory macrophage signature in CHIP (+) classical monocytes in severe COVID-19. **a**, GSEA plots for marker genes of inflammation clusters in CHIP (+) and CHIP (−), respectively. Genes are ordered based on log-fold changes between high inflammation cluster and low inflammation cluster. Normalized enrichment scores (NES) and FDR are presented for DEGs between M1-like and M0 macrophage^37^. **b**, the effect of SYK inhibitor to suppress CHIP (+) up-regulated genes. Comparison between combined scores of marker genes of high inflammation cluster for Kinase Perturbations from GEO down gene ontology library for inhibitors or knock out of SYK kinase terms. One-sided Paired t-test was performed (*: *P<* 0.05).

**Extended Data Figure 6.**
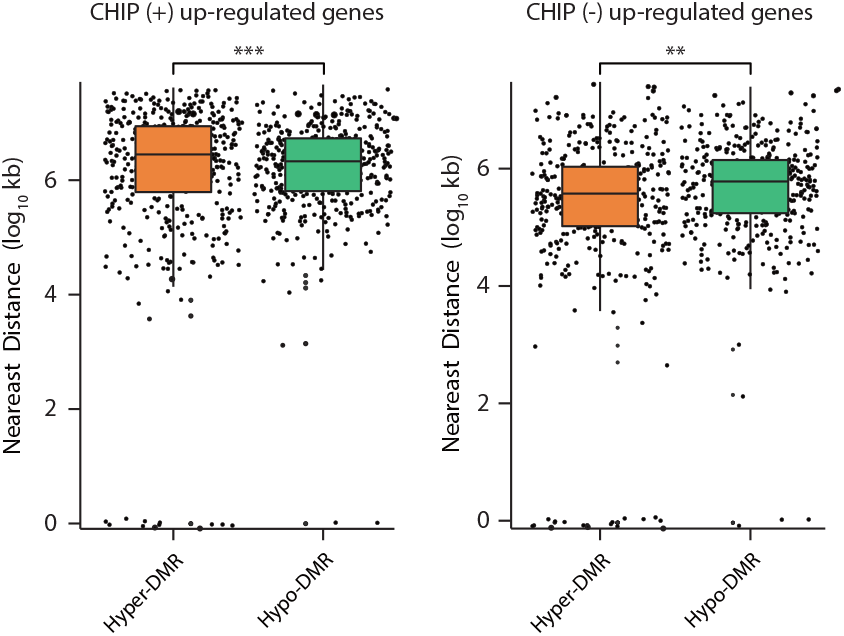
The distribution of the nearest distance between differently expressed gene promoters and DMRs. Boxplots of the nearest distance between differentially expressed genes (right: CHIP (+) up-regulated genes, left: CHIP (−) up-regulated genes) and DMRs. Hypo-DMRs were randomly selected to have the same sample size as hyper-DMRs. The box represents the interquartile range (IQR) and the whiskers correspond to the highest and lowest points within 1.5 × IQR. The color indicates the types of DMRs. Distance is represented as the log scale, with a 10kb resolution. For statistical significance test, two-sided Kolmogorov-Smirnov test were performed between hypo- and hyper-methylated regions (*P*<0.001, CHIP (+); *P*=3.46e-3, CHIP (−)).

## Supplementary Tables

**Supplementary Table 1. Baseline characteristics of COVID-19 patients in this study according to the presence or absence of clonal hematopoiesis of indeterminate potential**

**Supplementary Table 2. Clinical characteristics of cluster A2 when compared other cases in cluster A**

**Supplementary Table 3. Clinical information of patient used in scRNA seq**

**Supplementary Table 4. Data quality of scRNA seq**

**Supplementary Table 5. A list of DMRs**

